## Supplementary Information for "Heterogeneity of Rift Valley fever virus transmission potential across livestock hosts, quantified through a model-based analysis of host viral load and vector infection"

#### S.1 Within-host model of RVFV infection

##### S.1.1 Parameter estimation

The Deviance Information Criterion (DIC) is calculated as:

$$DIC = p_D + \overline{D(\theta)} \tag{S.1}$$

$$D(\theta) = -2.log(p(y|\theta)) + C \tag{S.2}$$

$$p_D = \overline{D(\theta)} - D(\bar{\theta}) \tag{S.3}$$

where $\theta$ is the vector of unknown parameters sampled, $p$ is the likelihood function and $y$ the data. We compute this DIC on the joint posterior distribution of 3 chains, each with the burn-in period removed.

For species heterogeneity, as no cattle nor goats died during the experiment, we based our comparison on surviving individuals. Therefore, we compute the DIC of a model fitted to the dataset comprising all surviving individuals, and compare it to the sum of DICs of models fitted to each species dataset separately. Similarly, for lambs, we compute the DIC of a model fitted to the whole lamb dataset, and compare it to the sum of DICs of models fitted to surviving and dying lambs separately.

Due to correlation between parameters, the value of $T_0$ could not be estimated within the MCMC procedure. Instead, we produced likelihood profiles by running the MCMC procedure for different values of $T_0$, for each animal group, and recorded the maximum log-likelihood (Figure S.1).

We then performed the Gelman-Rubin diagnostic to check for the common convergence of multiple chains (Figure S.2 for models selected by DIC). All multivariate potential scale reduction factors were <1.01 (Figure S.3). The joint posterior distributions are shown in Figure S.4.

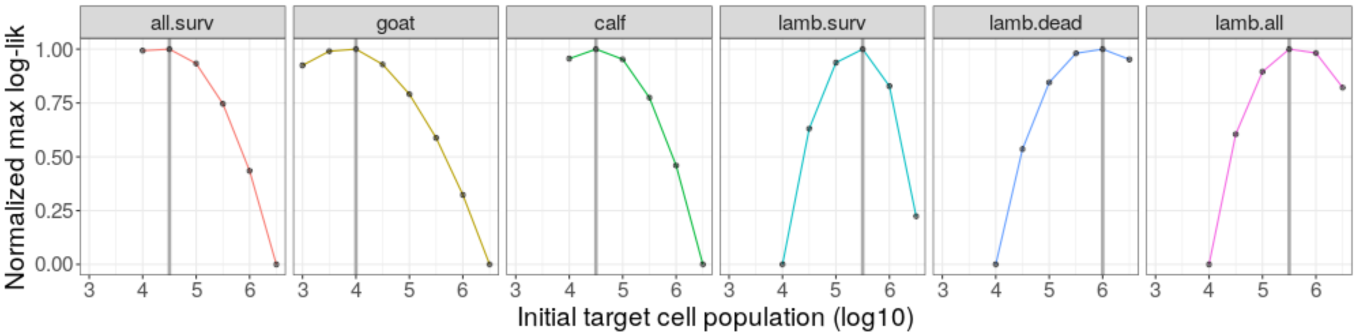


Figure S.1: Likelihood profiles to estimate $T_0$. Each point corresponds to the (normalized) maximum log-likelihood of an MCMC procedure run with a given $T_0$ value, for a given animal group (facet title). Vertical lines show the $T_0$ retained for further parameter estimation (chains of 100,000 iterations). Note that for lambs, assigning a common $T_0$ to individuals surviving and succumbing to the infection does not give the best fits so these groups were assigned different values.

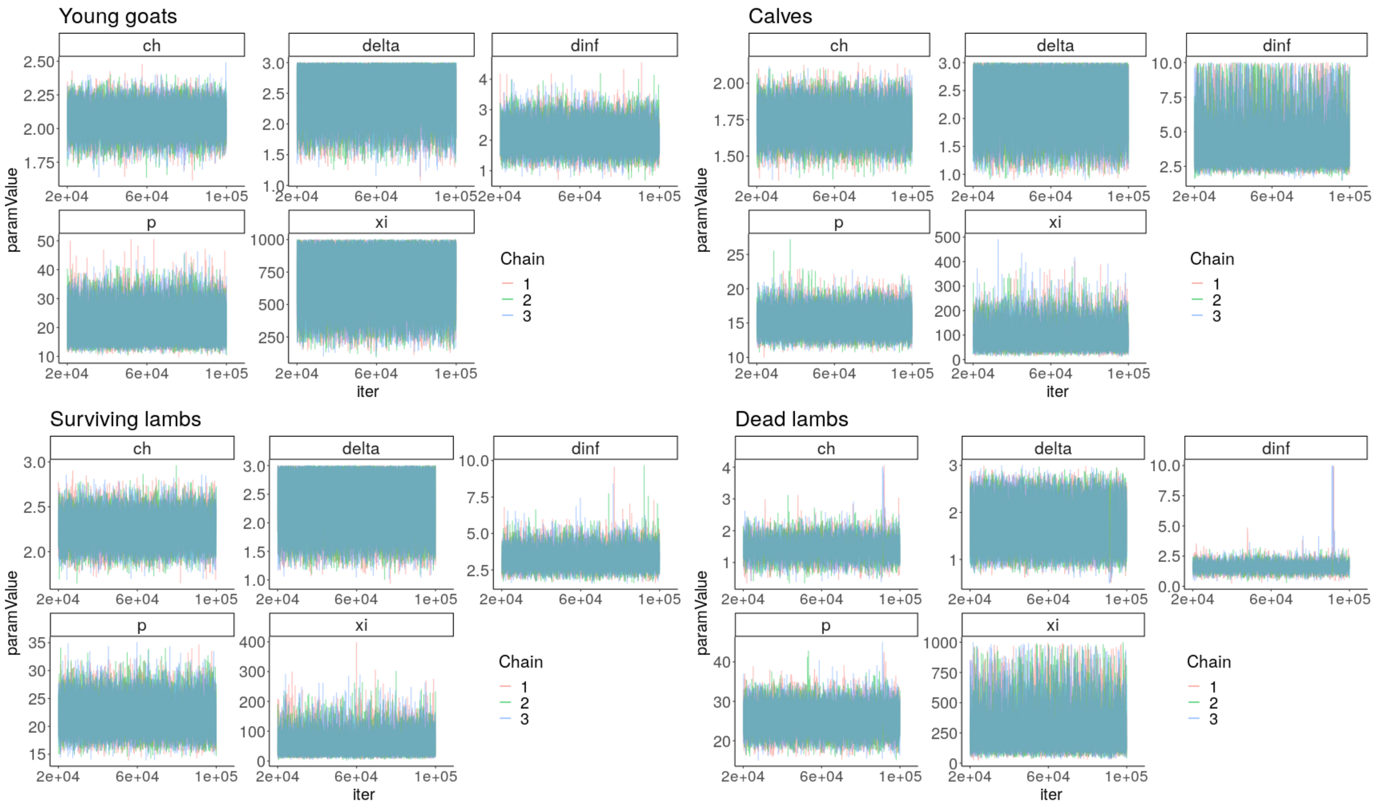


Figure S.2: Trace plots of selected models. Three chains, 100,000 iterations, 20,000 burn-in.

##### S.1.2 Outcome measures

According to Svensson, 2007:

For a population model, the expected time between a primary case and a secondary case is :

$$E[T_g] = E[L] + \frac{E[I^2]}{2E[I]}$$

For our within-host model, $I$ and $L$ now refer to the state of the cells instead of individuals, and we add $E[V_{inf}]$.

A property of the expected value is : $V[I] = E[I^2] - E[I]^2$

$$E[T_g] = E[L] + \frac{V[I] + E[I]^2}{2E[I]} + E[V_{inf}]$$

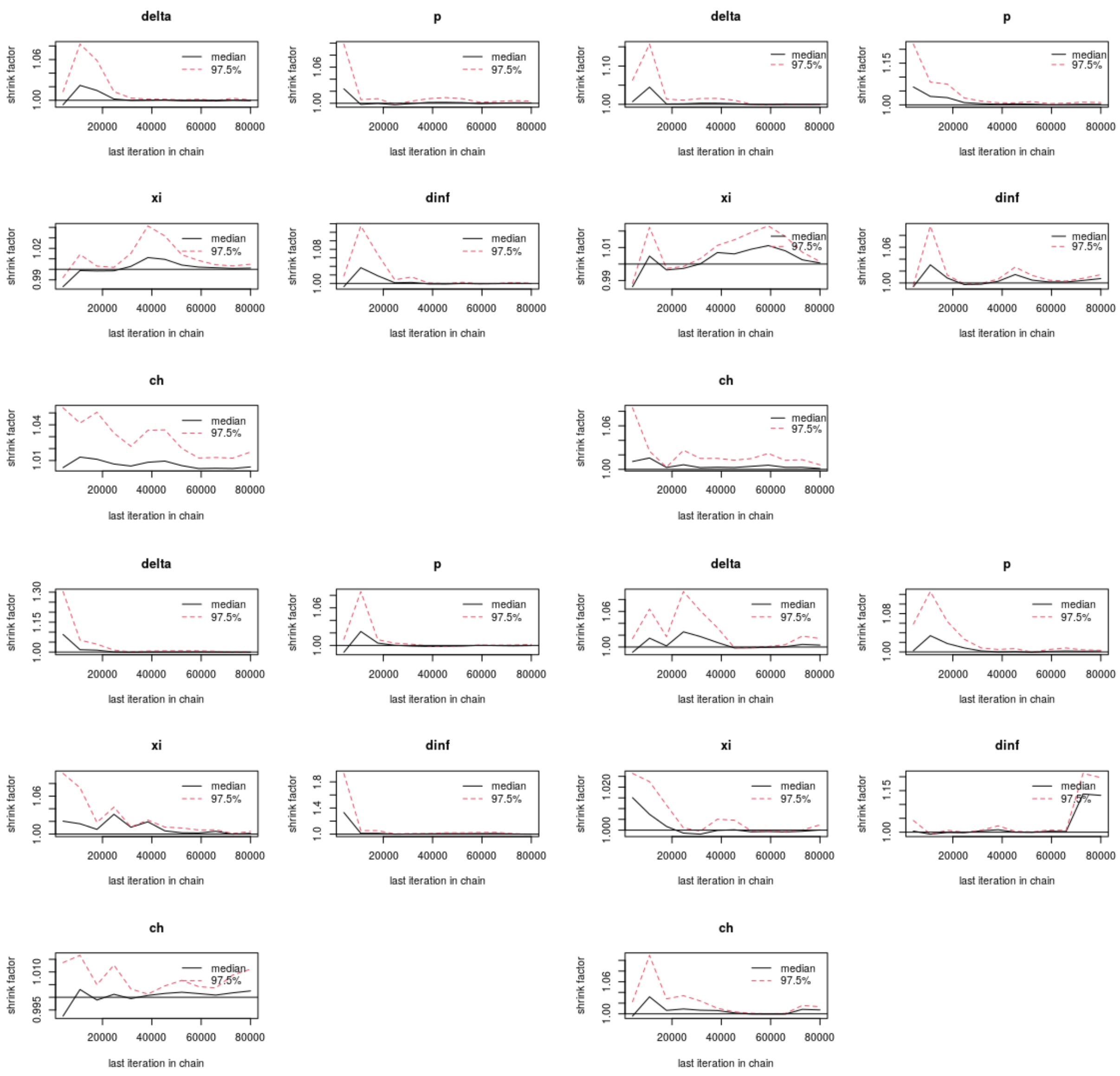


Figure S.3: Gelman diagnostic plots, per parameter for selected models. Top left : goats, top right : calves, bottom left : lambs which survived, bottom right : lambs which died.

for Erlang distributions, $V[I] = \dfrac{E[I]^2}{n}$, n : shape parameter

$$\frac{V[I] + E[I]^2}{2E[I]} = \frac{\frac{E[I]^2}{n} + E[I]^2}{2E[I]} = \frac{\frac{n+1}{n}E[I]^2}{2E[I]} = \frac{n+1}{2n}E[I]$$

$$E[T_g] = E[L] + \frac{V[I] + E[I]^2}{2E[I]} + E[Vinf] = \kappa^{-1} + \frac{n+1}{2n}\delta^{-1} + (c_h + d_{inf} + \sigma\beta T_0)^{-1}$$

### S.2 Dose-response relationship in RVFV mosquito vectors

#### S.2.1 Systematic review

A systematic review of the literature was performed in order to investigate the relationship between vertebrate host infectious titer at the moment of blood-feeding and infection of a vector. PubMed and Scopus were searched

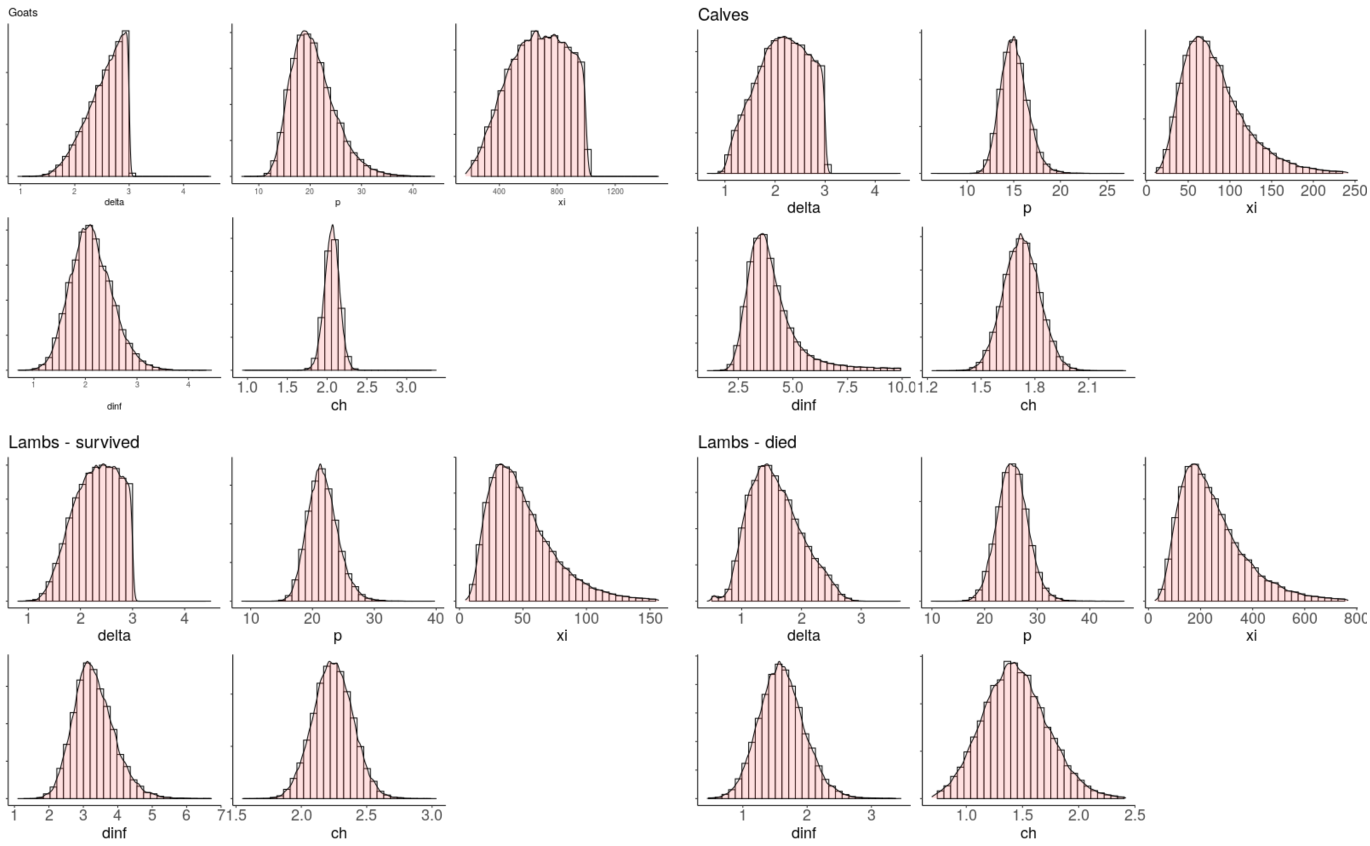


Figure S.4: Joint posterior distributions of parameters per selected model. Priors can be found in Table 2. $\delta$ is constrained to be inferior to $\kappa$ (fixed, $= 3$), and $c_h + d_{inf}$, as advised by Smith et al., 2010

on $27^{th}$ september 2021 with the following query:

```
((Vector*) OR (Mosquit*) OR (Aedes) OR (Culex) OR (Anopheles))
AND
((Transm*) OR ("vector competence") OR (Infect*))
AND
(("rift valley fever"))
AND
((Virol*) OR ("virus level") OR ("viral level") OR ("virus load") OR ("Viral load")
    OR (Virem*) OR (Viraem*))
AND
((Host*) OR (Ruminant*) OR (Sheep*) OR (Goat*) OR (Cow) OR (Lamb*) OR (Calve*) OR (Ewe)
    OR (Ewes) OR (Cows) OR (Buffalo*) OR (Camel*) OR (cattle) OR (rodent*) OR (hamster*))
```

This search provided 315 results in PubMed and 253 from Scopus, 357 of which were unique results. 316 articles were excluded at the title and abstract level, 23 at the full text level. The exclusion criteria (not mutually exclusive) for these steps were :

- no mention of RVFV (n = 26)
- no viral load values given (n = 215)
- no use of mosquito (n = 116)
- not written in English (n = 11)
- no presence of primary data (e.g number of mosquitoes, n = 51)
- full text not available (n = 9)
- review (n = 5)

Eventually, data from 18 articles was retrieved for further inspection. A descriptive analyses was performed to assess the diversity of protocols used. The 16 papers corresponded to 341 data points. Regarding hosts, 75% (n = 256) were hamster, 23% artificial feeding, and 2% lamb. Regarding mosquito vector used, 45% (n = 154) were *Culex* spp., 41% *Aedes* spp.. The rest was a mix of *Anopheles*, *Coquillettidia*, *Mansionia*, *Culiseta*, and *Psophora* spp.. Regarding RVFV strains, 73% were ZH501 (n = 250). The detail of other strains will be spared here, but none was used in more than 20 experiments.

We limited our quantitative analysis to experiments performed with *Aedes* and *Culex* spp., with strain ZH501, on hamsters. This was done because the number of experiments performed with other hosts and strains was too small to properly test for the effect of these variables. Hamsters have been shown to be good model hosts for RVFV infection (Scharton et al., 2015). ZH501 is a wild strain originally isolated from a human patient during the outbreak of 1977 in Egypt (Meegan, 1979).

##### S.2.2 Data description

Nine papers remained, corresponding to 185 data points (81 *Aedes*, 104 *Culex*). We first examined the distribution of timing of dissection, temperature and infectious titers in our dataset. The reporting of the duration between mosquito feeding and determination of its infection status was not always very precise. When a range was given, we used the middle point value ; when a lower limit was given, we used the limit plus 1 day. We applied the same rule for temperature and infectious titers, although the precision was usually better. The distribution of these 3 variables across experiments and for each genus is shown in Figure S.5. A Wilcox-test concluded that there was no significant difference between genus for these 3 variables.

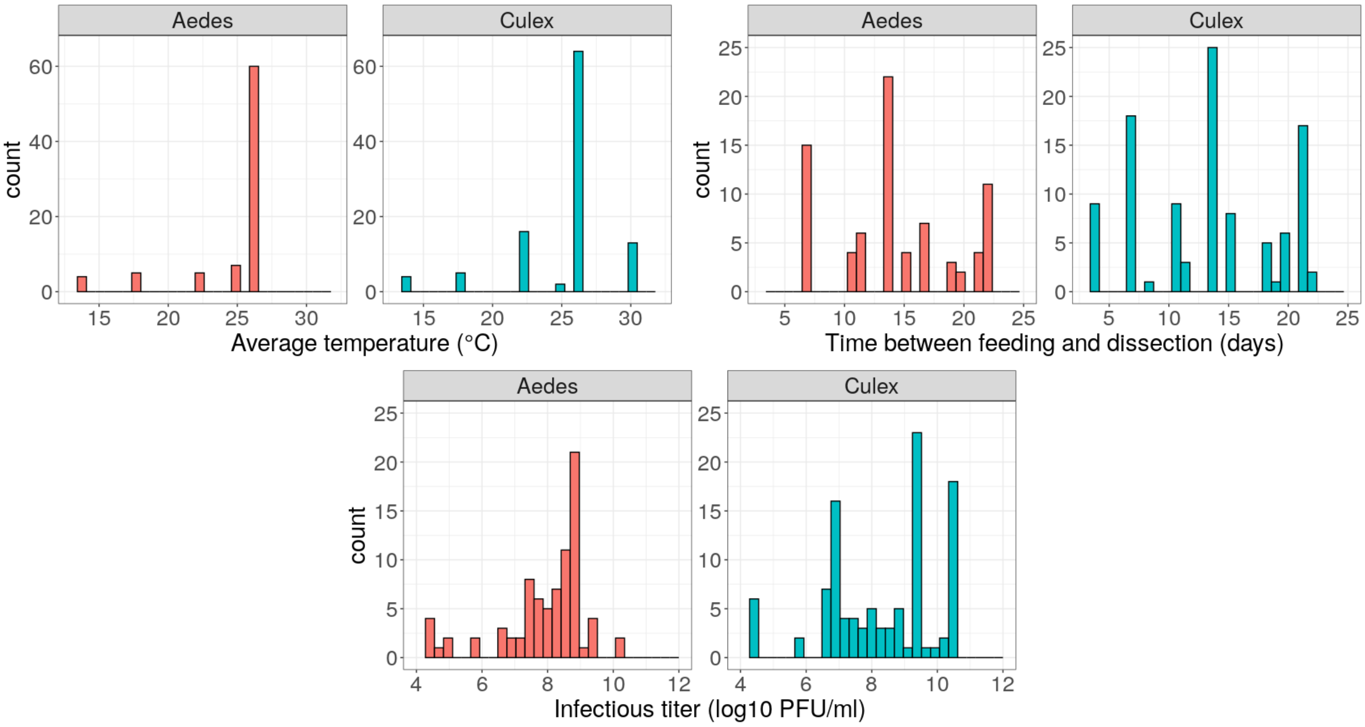


Figure S.5: Distribution of temperature, days post-exposure, and infectious titers, in experimental data retrieved from the systematic review, for *Aedes* and *Culex* spp. vectors.

```
            Wilcoxon rank sum test with continuity correction


data:  temp.aedes and temp.culex

W = 3847.5, p-value = 0.2275

alternative hypothesis: true location shift is not equal to 0

###

        Wilcoxon rank sum test with continuity correction


data:  dpf.aedes and dpf.culex

W = 4758, p-value = 0.1285

alternative hypothesis: true location shift is not equal to 0

###

        Wilcoxon rank sum test with continuity correction


data:  dose.aedes and dose.culex

W = 3536, p-value = 0.06089

alternative hypothesis: true location shift is not equal to 0
```

Note on units for infectious titers : In all experimental data retained for analysis, infectious titers were measured using plaque forming units (PFUs). This unit was kept for the fitting of dose-response equations. However, for the computation of livestock infectiousness, we converted PFUs into $TCID_{50}$ to be compatible with our time series of viral load. We used $V_{TCID_{50}} = V_{PFU} \times \frac{1}{0.69}$ which is consistent with 1 ml virus stock having half the number of PFUs as the $TCID_{50}$ (Canini et al. 2016).

##### S.2.3 Functional forms

We used a logistic regression model (Eq. S.4) to study the effect of temperature ($T$), duration between feeding of the mosquito and determination of its infection status ($dpf$) and infectious titer of the host it fed on ($dose$), on the probability to infect a mosquito. In our dataset, this probability is defined as the infection rate, i.e., the percentage of orally exposed mosquitoes that contained virus in their bodies (legs excluded). Indeed, dissemination rates (presence of virus in the head) showed strong variations, were measured on small sample sizes, and are likely to be influenced by other factors than infectious titers, out of the scope of this study. Transmission rates (presence of virus in the saliva) was only measured in 13% of our dataset.

$$F(T, dpf, dose) = \frac{1}{1 + e^{-(\beta_0 + \beta_1.T + \beta_2.dpf + \beta_3.dose)}} \quad \text{(S.4)}$$

We explored whether the infection data were over-dispersed using beta-binomial logistic regression. Based on AIC of models with or without overdispersal, we found that accounting for overdispersal was necessary (AIC for binomial model: 1509; AIC for beta-binomial model: 923). The beta-binomial model did not point towards a significant effect of temperature and number of days after feeding on infection rates. Dissection happened at least 4 days after blood feeding, which intuitively seems sufficient to infect a mosquito (before midgut barrier crossing).

```
                 Estimate  Std. Error z value     Pr(z)
beta0          -4.54506155  0.90396837 -5.0279 4.959e-07 ***
beta1           0.02605400  0.02419026  1.0770    0.2815
beta2          -0.00097779  0.01451044 -0.0674    0.9463
beta3           0.49687416  0.05892514  8.4323 < 2.2e-16 ***
overdispersion  4.14372201  0.56620726  7.3184 2.510e-13 ***
---
Signif. codes:  0 ‘***’ 0.001 ‘**’ 0.01 ‘*’ 0.05 ‘.’ 0.1 ‘ ’ 1
```

We then compared three functional forms (Eq. S.5-Eq. S.7) for the dose-response relationship, again using both a binomial and a beta-binomial likelihood for each. This was done on the whole dataset, *Aedes* and *Culex* spp. aggregated. Eq. S.5 is the logistic function, with the curve's maximum value fixed at 1 since we are dealing with probabilities. Eq. S.6 (Ferguson et al., 2015) was used to study vector competence for dengue when carrying *Wolbachia*. Eq. S.7 (Somvanshi and Venkatesh, 2013) is often used to model biological interactions that demonstrate sigmoidal response, in particular to capture the biomolecular interaction exhibiting cooperativity among two binding molecules.

$$Logistic(dose) = \frac{1}{1 + exp(-(\beta_0 + \beta_1.dose))} \tag{S.5}$$

$$Ferguson(dose) = 1 - exp(\frac{-dose}{\beta_0} + dose)^{\beta_1}) \tag{S.6}$$

$$Hill(dose) = \frac{dose^{\beta_0}}{\beta_1} + dose^{\beta_0} \tag{S.7}$$

Based on AIC, the Ferguson model best fitted the data. The AICs of all three functional forms supported the assumption that the data were betabinomially distributed.

```
                     AIC df
binom_Log      1532.8243  2
betabinom_Log   919.7607  3
binom_Ferg     1524.8107  2
betabinom_Ferg  918.8410  3
binom_Hill     1519.1427  2
betabinom_Hill  918.8682  3
```

We then fitted *Aedes* (n = 81) and *Culex* (n = 104) observations separately with Eq. S.6. The likelihood ratio test showed that the dose-response relationships associated with these two categories were significantly different ( Figure 4 in main text ).

```
> lrt(LL0 = betabinom_Ferg_CuAe@details$value,
+     LL1 = sum(betabinom_Ferg_Culex@details$value, betabinom_Ferg_Aedes@details$value),
+     df0 = 3, df1 = 6)
$L01
[1] 31.13315
$df
[1] 3
$‘p-value‘
[1] 7.969038e-07
```

Table S.1 shows the different species included in our dataset, with the associated number of observations. We explored species-specific curves for species that had sufficient data available at a large enough range of dosages. We ended up fitting a curve for *Cx. tarsalis*, *Cx. nigripalpus*, *Ae. vexans*, and *Ae. j. japonicus* (Figure S.6).

| *Aedes* spp. | | *Culex* spp. | |
|---|---|---|---|
| Species | n | Species | n |
| *Ae. taeniorhyncus* | 21 | ***Cx. tarsalis*** | 57 |
| ***Ae. vexans*** | 13 | ***Cx. nigripalpus*** | 15 |
| ***Ae. j. japonicus*** | 8 | *Cx. quinquefasciatus* | 10 |
| *Ae. infirmatus* | 8 | *Cx. pipiens* | 7 |
| *Ae. calceatus* | 6 | *Cx. salinarius* | 6 |
| *Ae. aegypti* | 4 | *Cx. erraticus* | 5 |
| *Ae. atlanticus* | 4 | *Cx. zombaensis* | 2 |
| *Ae. circumluteolus* | 3 | *Cx. annulirostris* | 1 |
| *Ae. dorsalis* | 3 | *Cx. erythrothoras* | 1 |
| *Ae. notoscriptus* | 3 | | |
| *Ae. vigilax* | 3 | | |
| *Ae. communis* | 1 | | |
| *Ae. fitchii* | 1 | | |
| *Ae. implicatus* | 1 | | |
| *Ae. sticticus* | 1 | | |
| *Ae. stimulans* | 1 | | |

Table S.1: Number of datapoints available per vector species, retrieved from the systematic review. *Ae. taeniorhyncus* was not selected as experiments performed on this species only included two distinct dosages.

#### S.3 Net infectiousness of RVFV livestock hosts

Figure S.7 shows the sensitivity of lambs' net infectiousness (NI) to the survival rate in the population. Indeed, NI is computed from 1000 dose-response curves combined with 1000 viral load curves, each of which is assigned to be of a surviving or dying type depending on the survival rate in the population. For viral load curves of dying individuals, we then apply a Weibull survival model directly computed from experimental times of death to truncate the viral load dynamics at the relevant timestep. We do not test the sensitivity of NI to those death times.

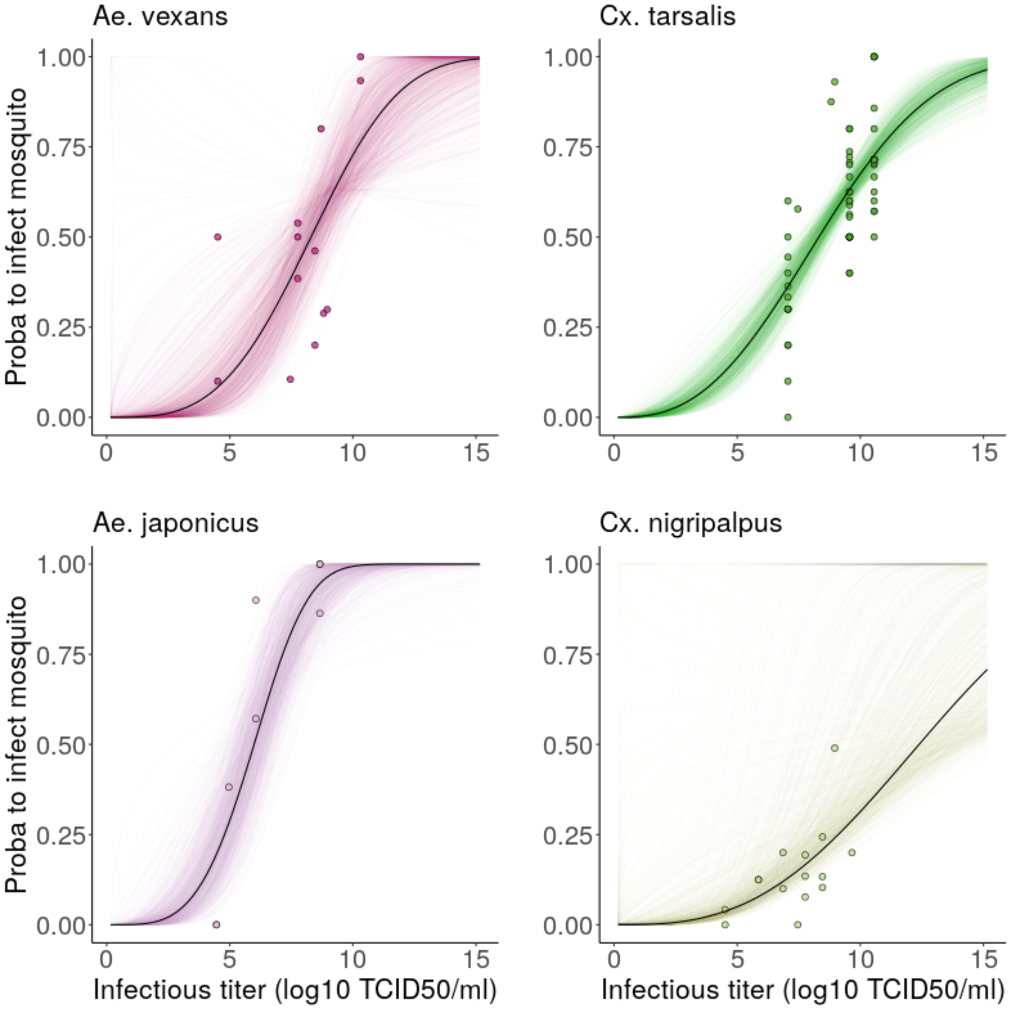


Figure S.6: **Species-specific dose-response curves.** Probability to infect a mosquito (y-axis) given feeding on a host with a certain infectious titer (x-axis). Points represent data gathered from literature review, where the probability is computed as the proportion of mosquitoes containing RVFV in their bodies (legs excluded), in a pool of fed mosquitoes (sample size not shown on plot for readability). Black lines show the dose-response relationship obtained by fitting Eq. S.6 with a betabinomial likelihood accounting for overdispersal in the data. Colored lines show the uncertainty around the fit : 1000 trajectories obtained by sampling in the variance-covariance matrix of the fitted model.

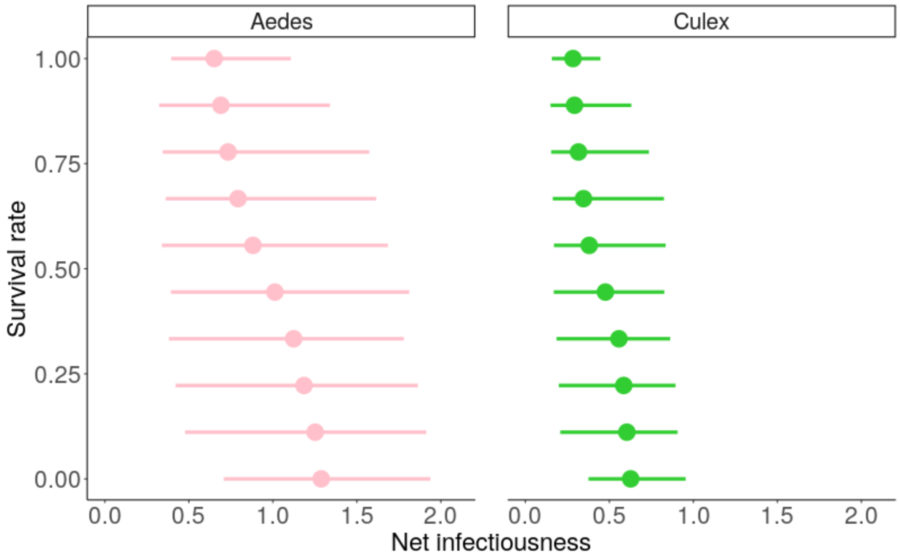


Figure S.7: Net infectiousness of an average lamb in relation with the expected survival rate in the population, for transmission to *Aedes* and *Culex* spp. vectors. Points show median, lines show highest density interval, computed from 1000 estimates.
